# Interactive effects of xylem and phloem damage under drought on plant water and carbon dynamics: Mimicking the impact of plant canker disease

**DOI:** 10.1101/2025.03.03.641225

**Authors:** Mohitul Hossain, Pieter Poot, Erik Veneklaas

## Abstract

- It is often stated that trees experiencing climate change may be more vulnerable to damaging effects of pest and disease, but experimental tests are still rare. In this study, we examined the separate and combined impact of experimentally applied xylem and phloem damage in an Australian Eucalypt species (*Corymbia calophylla)*, which is increasingly suffering from phloem and xylem damage by a canker disease caused by *Quambalaria coyrecup*.
- Cut treatments were applied to saplings under controlled conditions by removing xylem only (∼56%), phloem only (∼70%) or both in the main stem under well-watered and drought treatments.
- As expected, xylem damage reduced whole-plant conductance, stomatal conductance and photosynthesis, and hence growth. Phloem damage limited phloem transport, increased leaf non-structural carbohydrate concentrations (NSC), and reduced root NSC, photosynthesis and growth. Contrary to expectations, effects were larger in well-watered than droughted plants. The combined effects of damage to xylem and phloem were generally less than additive. Plants were remarkably resilient to significant loss of xylem and phloem, although xylem damage had a greater impact on most of the physiological parameters than phloem damage.
- These results reveal potential consequences of xylem-phloem dysfunction due to biotic attack, leading to whole tree mortality, in association with drought stress.

## Introduction

Rapid climate change, with increasing heat and drought events and associated pest and pathogen outbreaks, has been reported over the last few decades, and has been predicted to continue into the future with major consequences for forest productivity and survival (Allen *et al*., 2010; Matusick *et al*., 2013; Anderegg *et al*., 2015). A large number of tree mortality events in recent years across different ecosystems has increased interest in underlying mechanisms (McDowell *et al*., 2011; Adams *et al*., 2013; Hartmann *et al*., 2013; Mitchell *et al*., 2014; Aguadé *et al*., 2015). The theoretical framework of tree mortality proposed by McDowell *et al*. (2008) highlighted the close links between tree hydraulic function and carbon/energy economy. Trees may avoid drought-induced hydraulic failure by reducing transpiration, but this also leads to reduced photosynthesis which can in the long term cause carbon starvation. McDowell *et al*. (2008) moreover suggested that biotic agents may amplify drought-induced plant stress. One situation where this is likely, is where a pest or disease, such as wood borers, bark beetles or stem cankers, reduce the transport capacity of xylem or phloem, or both.

There is ample evidence that severe loss of xylem function plays a role in drought-related tree mortality in several species, but considerably less evidence for a role of carbon starvation (Adams *et al*., 2017). Concentrations of non-structural carbohydrates (NSC) decrease under drought in some species, but increase in others (Adams *et al*., 2017). Compared to plant hydraulics, the physiology of phloem function under drought is poorly known, but xylem and phloem transport are functionally linked. For example, declining water potentials in xylem require increased phloem osmolality to maintain phloem functionality (Sevanto, 2018). As sugars contribute significantly to osmolality, both the concentrations and transport of sugars in phloem are affected by low water potentials. Drought stress therefore will affect source-sink relationships, and sinks such as fine roots, which are far away from leaves and of vital importance for water uptake, may experience reduced growth and functionality.

Despite the tight hydraulic connection between xylem and phloem, empirical studies of their interactive effects on plants are very rare (Zwieniecki *et al*., 2004). Xylem and phloem physiology are often studied separately, and most phloem physiology studies are conducted using well-watered plants (De Schepper *et al*., 2010; Asao & Ryan, 2015), so that very little is know about the impact of impeded phloem transport under drought stress (Sevanto, 2014; Oberhuber *et al*., 2017). Experimental manipulation of xylem and phloem functionality has a long history, but has often involved rather extreme treatments such as overlapping sawcuts of sapwood (Greenidge, 1955; Postlethwait & Rogers, 1958; Schulte & Costa, 2010), and complete bark girdling (Williams *et al*., 2000; Li *et al*., 2003; Oberhuber *et al*., 2017). Abiotic or biotic factors are more likely to cause partial damage to xylem and/or phloem functionality. In the present study we investigate the effect of experimental damage to xylem, phloem or both, under well-watered as well as water-limited conditions.

*Corymbia calophylla* is a keystone eucalypt species in Western Australia, which has increasingly been suffering from stem canker infestations by the fungal pathogen *Quambalaria coyrecup* (Paap *et al*., 2008). Currently, the disease is widespread across *Corymbia calophylla*’s distribution range in trees of all age classes, causing mortality in extreme cases (Paap *et al*., 2017). This canker pathogen can attack and spread on branches and main stems, damaging cambium and vascular tissues (phloem and xylem) of host plants. From an infection point in the bark, the canker grows inwards and sidewards, creating a lesion of increasing size in the bark and sapwood, and causing loss of stem hydraulic conductance (Hossain *et al*., 2018). It is not known if the canker affects xylem more than phloem. Growth conditions likely influence the relative impact of xylem versus phloem damage: xylem function may be more critical for tolerance of drought, and phloem function may be more critical for growth potential in more favourable conditions. This study was designed to determine the separate and combined impact of xylem-phloem dysfunction, as caused by canker disease, by experimentally severing stem vascular tissues. In this study, we tested the following hypotheses: (1) Xylem loss will reduce whole-plant hydraulic conductance causing partial stomatal closure, leading to reduced photosynthesis and plant growth; (2) Phloem loss will increase NSC in leaves and reduce NSC in roots by disrupting phloem turgor pressure gradients; the increased sugar concentration in leaves will down-regulate photosynthesis; (3) During drought, xylem loss will limit leaf function more than in well-watered conditions, whereas the impact of phloem loss will be larger in well-watered plants.

## Materials and Methods

### Plant materials and growth condition

*Corymbia calophylla* plants were established in small pots (150 ml) from seeds collected from a native marri stand in Mossop, Western Australia (34°71’190” S 116°46’956” E), and after eight months, plants were transplanted to 1-litre pots filled with potting mix and river sand (1:1.5), and grown in a shadehouse. Two months before the experiment started, 120 healthy similar sized plants (average diameter at 5 cm above soil: 5.27 ± 0.06 mm, average total stem length: 0.218 ± 0.06 m) were transferred to PVC cylinders (450 mm height, 150 mm diameter) filled with 10.4 kg of soil mix. The cylinders had holes in the bottom to facilitate drainage, and were covered with a 1 mm nylon mesh to prevent soil loss. The soil surface was covered by a layer of gravel (3.8 cm depth) to reduce soil evaporation. Prior to the experiment, plants were moved to a greenhouse equipped with UV-transmissible glass. Plants were randomly allocated to five different stem cut treatments with two levels of watering (droughted and well-watered) (details below), and placed on five different benches that had an equal number of plants from every treatment combination. During a two-month acclimation period, plants were watered regularly to the point of excess water draining through the drainage holes. In the next section, watering treatments were explained in detail. The glasshouse was temperature controlled by evaporative cooling and a shade screen during the summer months. Air temperature ranged from 15°C to 33°C, with a mean of 23°C.

### Watering treatments

Well-watered plants were maintained at 12% gravimetric soil water content (SWC) throughout the experiment by daily rewatering PVC cylinders with the amount of water lost during the previous measurement interval, as determined by pot weighing. The drought treatment was commenced by cessation of watering until the pot SWC reached 5%, after which plants were maintained at that level by watering every 2-3 days throughout the experiment. Every pot was weighed to the nearest 0.1 g between 8:00 and 10:00 am to determine water loss.

### Xylem and phloem damage treatment

After a month of watering treatment, plants of each treatment were allocated to different xylem/phloem cut treatments (Fig. 1). For the phloem cut treatment, a 6 mm strip of bark was removed using a sharp blade for 70% of the stem perimeter. For the xylem cut treatment, 65-70% of xylem was removed using a flexible shaft rotary tool (Foredom, Bethel CT, USA) with a 2 mm burr, entering through a small hole in the bark, and moving sideways and inward to create a round-shaped cavity. In total, five treatments were established: Control, i.e. no cuts (C); Phloem cut (P); Xylem cut (X); Xylem and phloem cut (XP); and an additional control for X, having only the small bark entry hole (XC) as this caused a small amount of damage to phloem tissue (Fig. 1). After making the cut on phloem or xylem, the wound was blocked with silicon gel (Selleys, DuluxGroup Pty Ltd, Australia) and wrapped using Parafilm (Pechiney Plastic Packaging, Menasha, WI, USA), to avoid infection, desiccation and leakage. To quantify the percentage of sapwood/xylem cut, at the harvest, the stem was sliced at the cut treatment point and photographed under a compound microscope equipped with a digital camera (Nikon SMZ800, Digital Sight DS-Fi2, Nikon Corporation, Japan). The photographs were used to quantify the percentage of sapwood cut using ImageJ v 1.48 (National Institutes of Health, Bethesda, MD, USA). The average percentage of sapwood removed varied from 48% to 62% (average of 53.9 ± 1.1%) in well-watered plants, and from 51% to 66% (average of 57.8 ± 1.3%) in droughted plants, with no significant differences between watering treatments (*P* = 0.13). Each of the five treatments had 10 replicate plants in both well-watered and droughted conditions.

**Figure 1:**
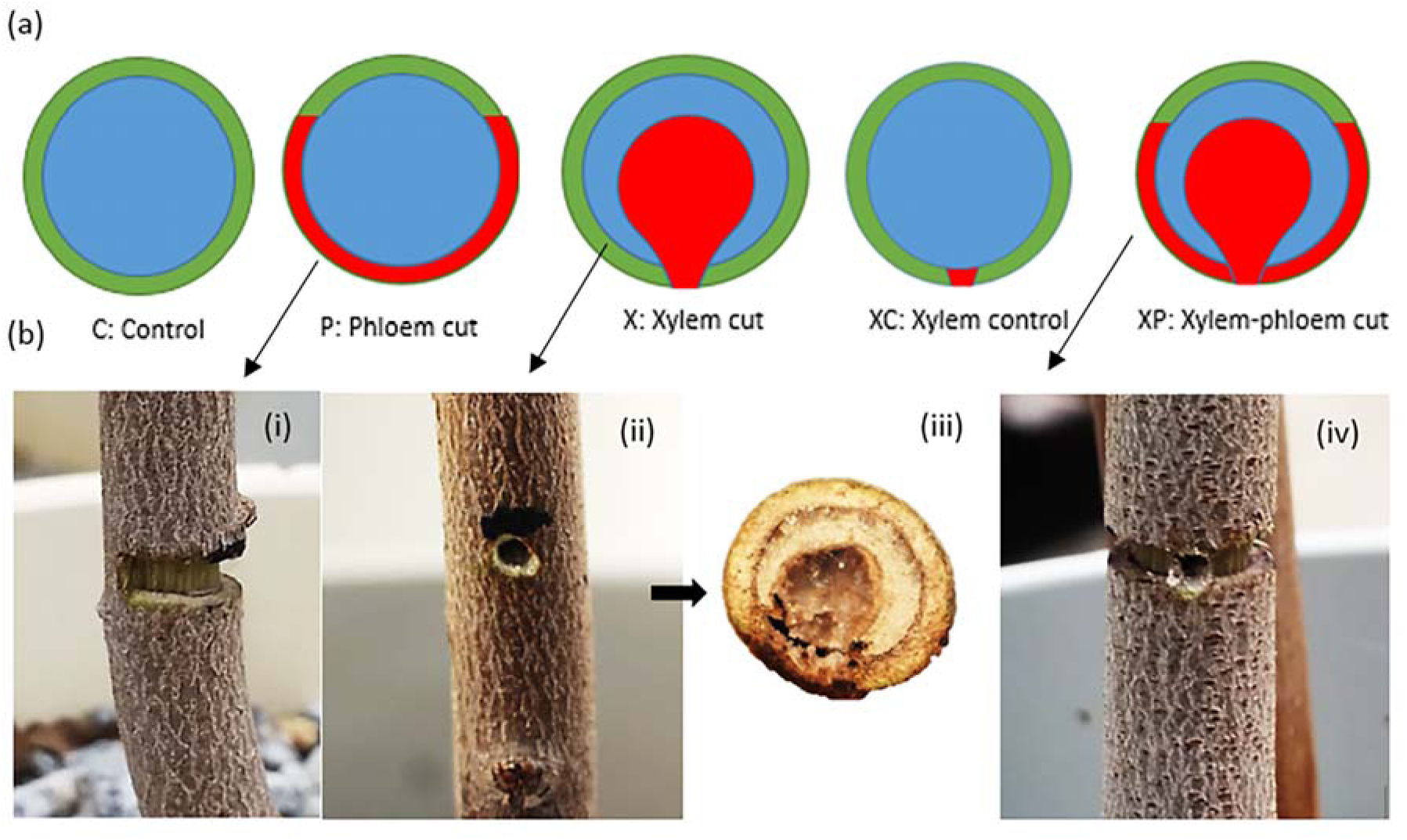
(a) Schematic stem cross-sections illustrating the five different xylem/phloem cut treatments; (b) Photographs of treated stems – i: Phloem cut; ii: Xylem cut; iii: Cross-section of Xylem cut; iv: Xylem-phloem cut.

### Growth measurement

Stem diameter (at the point of the cut, 50 mm below and 50 mm above), plant height and total stem length (height plus all branch lengths) were measured just before the cut treatment and at the final harvest, four months after xylem and phloem cut treatments were applied. At the end of the experiment, aboveground and belowground biomass were determined by measuring dry weights of stems and leaves, and roots (after four days in a 70°C oven). Before drying, roots were well rinsed under running water above a sieve to remove soil particles. Leaf area of the leaves from two branches of each plant was determined with a leaf area meter (Li-cor model LI-3000, LI-COR Inc., Lincoln, NE, USA), after which they were oven dried. These data were used to estimate whole plant leaf area from whole plant leaf dry weight.

### Water relations

Prior to the cut treatment, only midday leaf water potential was measured with a pressure chamber (Model 600, PMS, Albany, OR, USA), whereas at final harvest both predawn and midday leaf water potential were measured. Pre-dawn water potential measurements were carried out before sunrise between 4:00 and 5:00 am and midday water potential between 12:30 and 13:30 pm in the glasshouse. To estimate whole-plant conductance, all plants were placed in a growth chamber with a constant temperature (25°C) for two days at ∼115 days post cut treatment (i.e. one week before the final harvest). Well-watered plants were watered to their set SWC level in the late afternoon one day before the measurement, whereas droughted plants were watered in the late afternoon two days before the measurement to ensure an appropriate level of drought stress. To prevent soil evaporation, pots were sealed with plastic around the base of the stem. Whole-plant transpiration was determined by weighing (0.01 g accuracy) each pot 1 h before and 1 h after the midday water potential measurement. Transpiration was calculated from the water lost over this period. Midday leaf water potential was determined on one leaf per plant using a pressure chamber (Model 600, PMS, Albany, OR, USA). Immediately after the reweighing, all pots were moved to a dark, humid room and left for a minimum of 20 hours to ensure equilibrium between plant and soil water potential. During the following morning, predawn water potential was measured on one leaf per plant. Whole-plant conductance (g s^-1^ MPa^-1^ m^-2^) was calculated by dividing whole-plant transpiration (g s^-1^ m^-2^ leaf area) by the difference between midday and predawn water potential.

### Gas exchange

Leaf gas exchange was measured in the morning (8:00 – 10:30 am) in the glasshouse at ∼115 days post cut treatment using a LI-COR 6400 with a red-blue light source (LI-COR Inc., Lincoln, NE, USA) to assess the effects of drought and phloem-xylem damage. Gas exchange measurements on droughted plants were conducted on the second day after watering, to avoid possible effects of recent watering. A sun-exposed mature leaf was chosen from each replicate plant. For all measurements, we used a reference CO_2_ concentration of 400 μmol CO_2_ mol^−1^ and light intensity of 1500 μmol m^-2^ s^-1^. Leaf temperature was set at 25°C and relative humidity was 50 ± 10% during the measurements. Flow rates were set at 500 µmol s^−1^ for well-watered plants and 300 µmol s^−1^ for droughted plants.

CO_2_ response curves (A-Ci curves) were measured on 5-7 randomly chosen plants from each treatment combination, except the xylem control treatments (WXC and DXC) as no significant difference in photosynthesis was observed compared to controls (WC and DC), presumably due to the very limited damage of phloem. A-C_i_ curves were initiated after equilibrating the leaves for approximately 10 min at 400 μmol mol^−1^ CO_2_, 1500 µmol m^-2^-s^-1^ PAR and with leaf temperatures maintained at 25°C. The measurement was continued with a stepwise reduction in CO_2_ concentrations to sub-ambient (∼ 40 μmol mol^−1^) and then increased up to 1800 μmol mol^−1^, in a total of 12 steps. Leaves were maintained at each CO_2_ level for 120 to 240 seconds before being measured.

Stem damage may reduce sugar export from leaves, leading to increased sugar concentration, which may cause limitations in phosphate availability for the Calvin cycle (Lambers *et al*., 2008). The resulting feedback inhibition of photosynthesis can be detected by comparing A-Ci curves measured under ambient and low O_2_ (photorespiratory-repressing) conditions. According to biochemical modelling (Farquhar *et al*., 1980), light-saturated photosynthesis A_sat_ is stimulated under low O_2_ compared to ambient O_2_ in both the carboxylation limited region (V_cmax_) and the RuBP regeneration region (J_max_). However, plants in feedback-inhibited conditions are likely to show O_2_ insensitivity (no increase of A_sat_) or reverse sensitivity (reduction of A_sat_) to low O_2_. Therefore, we measured A-C_i_ curves at low oxygen (2 kPa) along with ambient oxygen (21 kPa; described above), as described in Ellsworth et al. (2015) to assess feedback inhibition of photosynthesis. The analysis of the A-C_i_ curves was done according to Sharkey (2016).

### Non-structural carbohydrates

Leaf, stem (0-50 mm above the cut) and root samples were collected at harvest and were kept at -20°C. Samples were freeze-dried (VirTis Benchtop K, SP Industries Inc., Warminster, PA, USA) and ground with a Geno/Grinder (SPEX SamplePrep 2010, Metuchen, NJ, USA). Fifty mg of ground sample material was extracted with 500 μL of 80% (v/v) ethanol in a hot water bath (30 min, 80°C). Samples were centrifuged for 15 min at 15,339 *g* (Biofuge 13, Heraeus Instruments, Hanau, Germany) after which the supernatant was transferred into Eppendorf tubes. The pellet was then resuspended in 500 μL of 80% (v/v) ethanol for a second and third extraction. Supernatants of the extractions were combined and 400 μL were transferred into HPLC tubes. The extracts were analysed for glucose, fructose and sucrose contents using HPLC (Chow & Landhäusser, 2004). The pellet was dried in a fume hood before adding 600 μL of enzyme mix. The enzyme mix consisted of 2000 μL of 10% amyloglucosidase solution and 20 μL of 10% α-amylase (Roche Diagnostics, Indianapolis, USA) in 50 mL of acetate buffer (50 mM). Samples were then incubated for 36 hours at 37°C and subsequently centrifuged for 10 min at 15,339 *g*. The supernatant was transferred into HPLC tubes for analysis. The starch concentration (g g^-1^) of the samples was calculated by multiplying the glucose concentration with 0.9. A Waters 717 autosampler and 600E dual head pump (Waters Corp., Milford, USA) were used in combination with an evaporative light scattering detector (Grace, Columbia, USA). Separation was performed on a Grace Prevail™ ES Carbohydrate column (250 mm x 4.6 mm) with 5 μm particle size at 30°C. The mobile phase consisted of 75% acetonitrile in milli-Q water (at a flow rate of 1 mL min^-1^).

### Statistical analysis

For statistical evaluation of the data, we used the statistical software package R (v3.4.1; R Foundation for Statistical Computing, Vienna, Austria). Two-way analysis of variance (ANOVA; function ‘aov’) was used to assess the effects of cut and water treatments on plant growth, water relations, gas exchange parameters and carbohydrates. Further, a Tukey–Kramer test (*P* < 0.05) was used to identify the significant differences across the treatments when significant effects were detected. To evaluate the relationship of A_sat_ with g_s_ and total soluble sugars across the cut treatment and watering treatments, we used a linear model (function ‘lm’). Further, we calculated a V_cmax_ ratio and a J_max_ ratio, measured at low versus ambient partial pressure of oxygen, and used a one sample t-test to assess if it was different from unity, indicating a significant effect of oxygen partial pressure. Before the analysis, data were log-transformed if necessary to correct for non-normality and heteroscedasticity.

## Results

### Plant growth

At the onset of the watering treatment, plants allocated to different treatments in the experiment were uniform in size (mean diameter at 50 mm above-cut point of 7.30 ± 0.07 mm; mean total stem length of 0.265 ± 0.006 m; P = 0.43 and *P* = 0.68 respectively for ANOVAs comparing treatments). The drought treatment significantly reduced growth compared to the well-watered treatment in all treatment combinations (P < 0.001; Fig. 2). Plants with xylem and/or phloem cuts had lower relative stem (length) growth rate than non-cut controls in both watering regimes, but this was only statistically significant for the well-watered plants (Fig. **2a**). The reduction of relative stem growth rate in these well-watered treatments, compared to the WC control, was 26% in the phloem cut (P = 0.73), 57% in the xylem cut (P = 0.04) and 65% in the xylem-phloem cut plants (P = 0.03; Fig. **2a**). Relative stem diameter growth above and below the cut was significantly lower in droughted treatments (P < 0.001), but only above-cut diameter growth differed between cut treatments, and only in the well-watered treatments (P = 0.02; Table S1). This was due to increased above-cut diameter growth in treatments involving phloem cuts: mean growth for WP and WXP was 4.64 mm, whereas mean growth of WC, WXC and WX was 3.52 mm. Similar to the results for stem growth rates, total plant biomass was only significantly decreased in well-watered xylem-cut (21%; P = 0.02) and xylem-phloem cut plants (26%; P = 0.002; Fig. **2b**), compared to the WC control. Root, stem and leaf fractions showed the same trends as total biomass (Fig. **2b**), however, root:shoot ratio was on average 22% higher in plants subjected to drought compared to well-watered plants (0.77 vs. 0.63; P < 0.001; Fig. S1). Well-watered phloem cut plants had the lowest root:shoot ratio.

**Figure 2:**
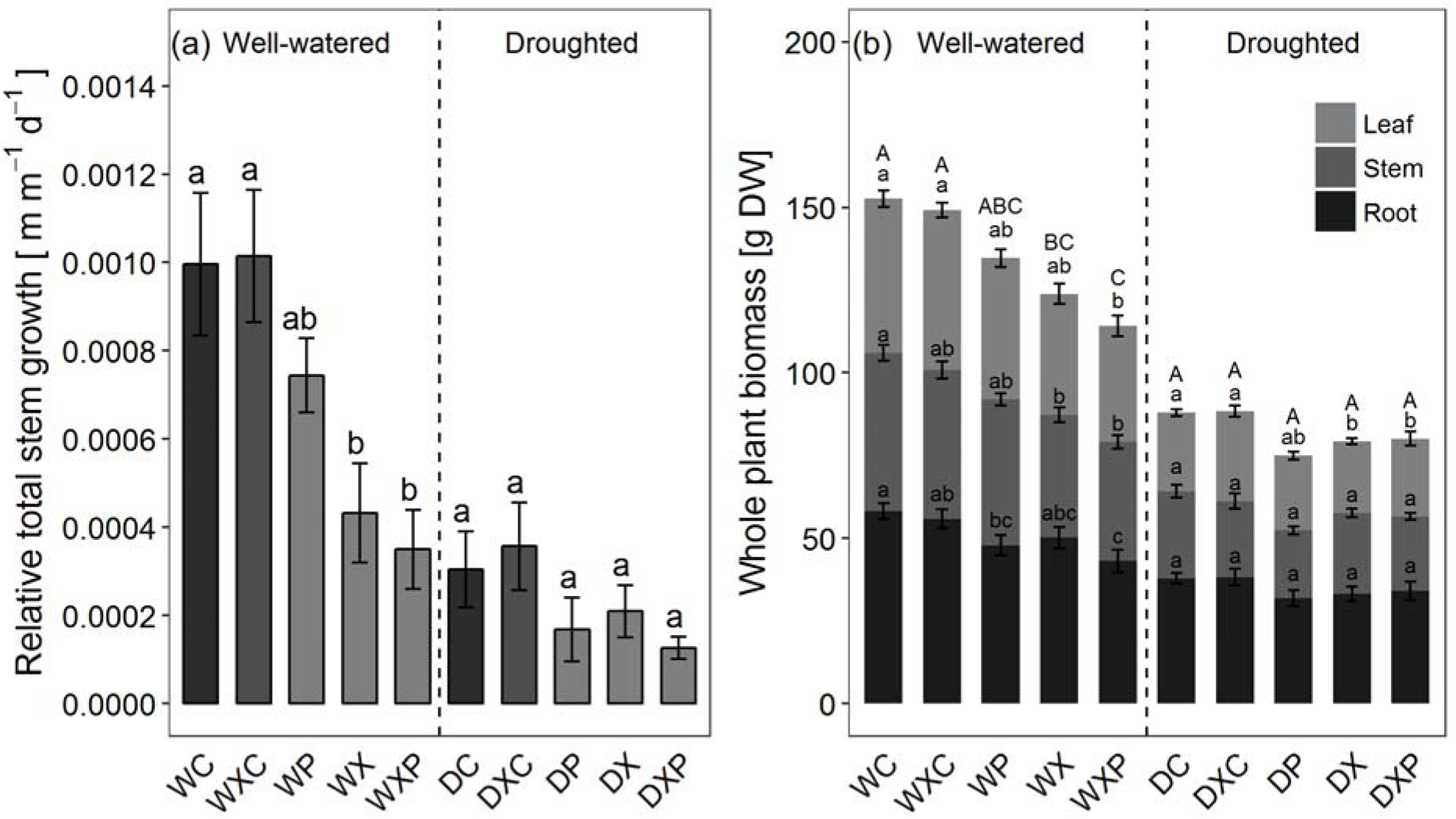
(a) Relative growth rate of total stem length over the experimental period and (b) whole-plant dry biomass at harvest. Different letters indicate significant (*P* < 0.05, Tukey– Kramer test) differences between cut treatments within each water treatment. Lower-case letters indicate significant (*P* < 0.05, Tukey–Kramer test) differences in relative total stem growth rate in Fig. 2a and in different plant parts (leaf, stem and root) in Fig. 2b; capital letters indicate differences in whole plant biomass in Fig. 2b. Treatments: WC = Well-watered control, WXC = Well-watered xylem control, WP = Well-watered phloem cut, WX = Well-watered xylem cut, WXP = Well-watered xylem-phloem cut, DC = Drought control, DXC = Drought xylem control, DP = Drought phloem cut, DX = Drought xylem cut, DXP = Drought xylem-phloem cut.

### Water relations

Throughout the experiment, droughted plants had lower midday leaf water potential (-1.8 to -2.6 MPa) than well-watered plants (-0.8 to -1.4 MPa) (Fig. **3a**). At final harvest, both predawn and midday water potential were significantly lower in droughted plants than well-watered plants (P < 0.001), but midday water potential were only different among the cut treatments in the well-watered plants (P > 0.05). Plants that had xylem cuts in the well-watered treatment (WX and WXP) had significantly lower midday water potential than well-watered controls (WC, WXC) (Fig. **3a**). Droughted plants also had lower whole-plant hydraulic conductance, K_p_, than well-watered plants (P < 0.001; Table S1). Similar to the leaf water potentials, K_p_ did not differ among the cut treatments for droughted plants, whereas it did for well-watered plants. In well-watered plants, the strongest reduction of K_p_ compared to control plants was observed in xylem-phloem cut plants (42% reduction; P = 0.004), followed by xylem cut (38% reduction; P = 0.008) and phloem cut plants (16% reduction; P = 0.78; Fig. **3b**).

**Figure 3:**
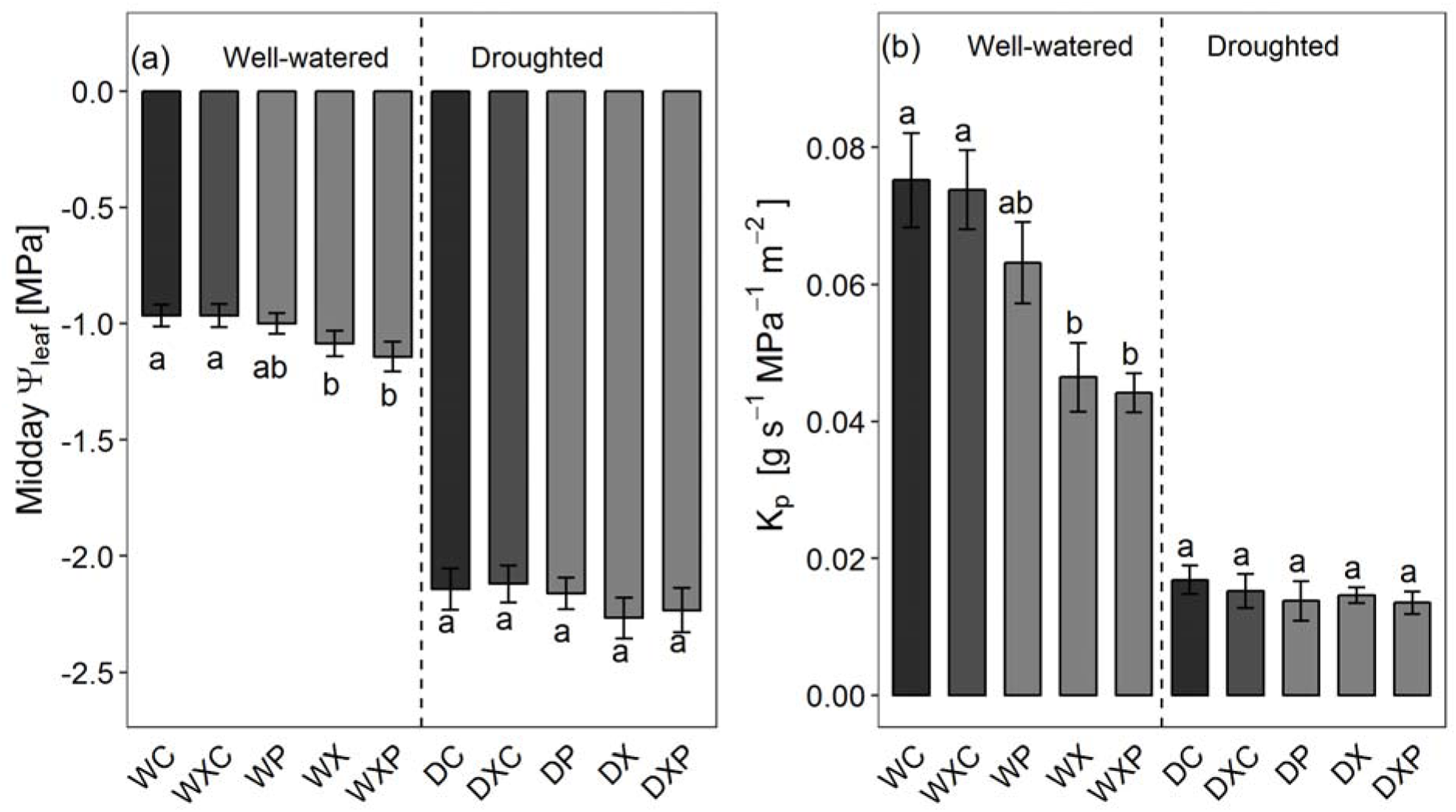
(a) Midday water potential and (b) whole-plant hydraulic conductance (K_p_) at harvest. Different letters indicate significant differences (*P* < 0.05, Tukey–Kramer test) between cut treatments within each water treatment. Treatments: WC = Well-watered control, WXC = Well-watered xylem control, WP = Well-watered phloem cut, WX = Well-watered xylem cut, WXP = Well-watered xylem-phloem cut, DC = Drought control, DXC = Drought xylem control, DP = Drought phloem cut, DX = Drought xylem cut, DXP = Drought xylem-phloem cut.

### Gas exchange

Before the final harvest, droughted plants had lower photosynthetic rates (A_sat_, 1.0–7.1 µmol CO_2_ m^-2^ s^-1^) than well-watered plants (6.6–16.6 µmol CO_2_ m^-2^ s^-1^) (P < 0.001; Fig. 4; Table S1). In the well-watered treatment, photosynthetic rate was significantly reduced by xylem damage (22% reduction compared to control; P = 0.01) and xylem-phloem damage (28% reduction; *P* < 0.01), and marginally reduced by phloem damage (∼12% reduction; P = 0.35) (Fig. 4). A similar trend was observed in droughted plants (Fig. 4). Across all the treatments, there was a tight correlation between photosynthesis and stomatal conductance (P < 0.001; R^2^ = 0.79). While reduced photosynthesis was largely explained by stomatal closure, A-C_i_ curve parameters also suggest a modest decline of V_cmax_ and J_max_ (Table 1). This reduction in V_cmax_ compared to that of well-watered control plants was statistically significant for the well-watered xylem cut (17%) and droughted xylem cut (34%) plants (Table 1). Under low O_2_, V_cmax_ and J_max_ were not significantly affected compared to ambient O_2_, except for a significant reduction in well-watered phloem-damaged plants (Table 1).

**Figure 4:**
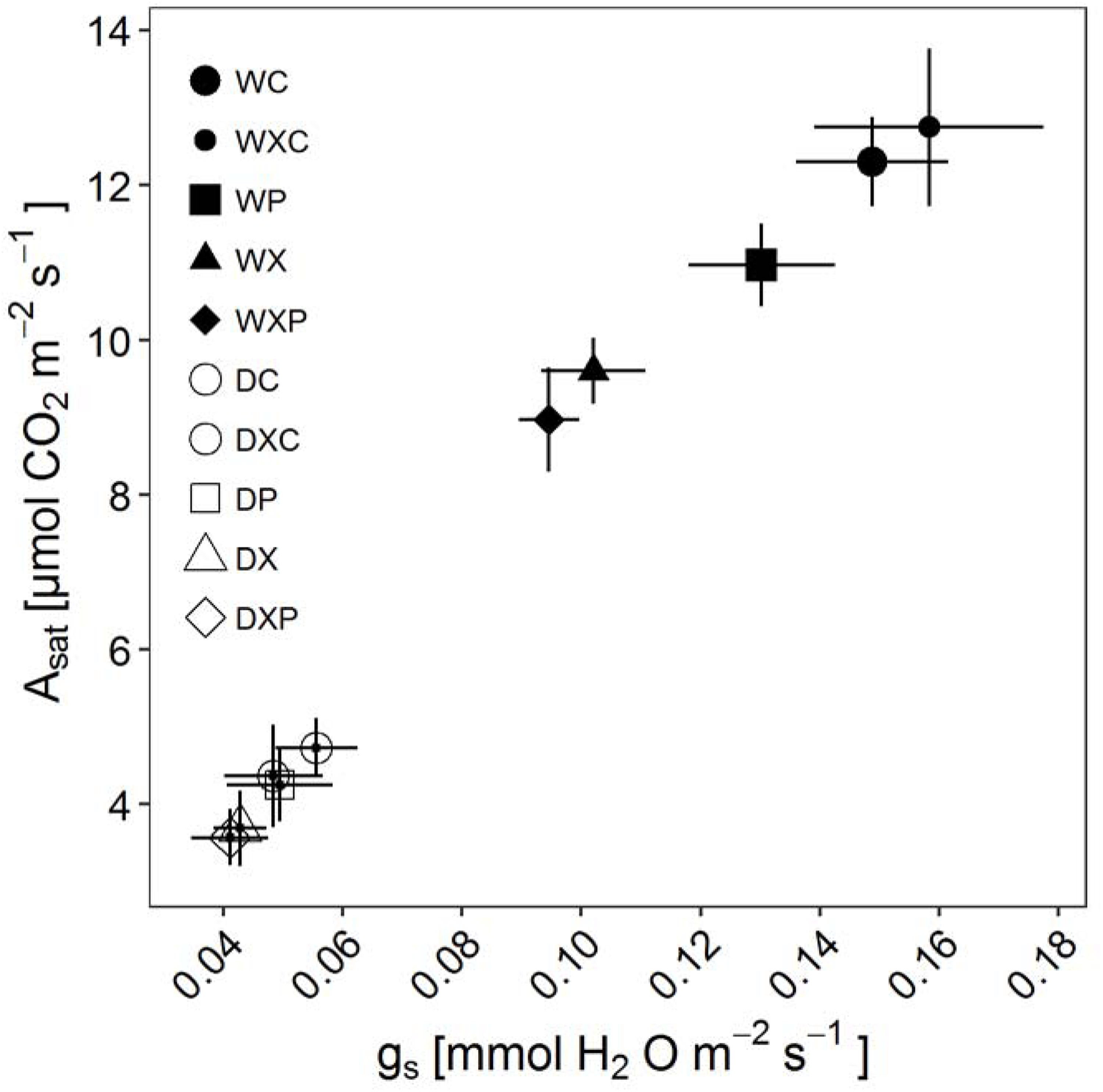
Net photosynthetic rate at saturating light intensity (A_sat_) as a function of stomatal conductance (g_s_). Treatments: WC = Well-watered control, WXC = Well-watered xylem control, WP = Well-watered phloem cut, WX = Well-watered xylem cut, WXP = Well-watered xylem-phloem cut, DC = Drought control, DXC = Drought xylem control, DP = Drought phloem cut, DX = Drought xylem cut, DXP = Drought xylem-phloem cut.

**Table 1:** V_cmax_ and J_max_ measured at ambient oxygen partial pressure, and the ratio of both parameters when measured at low and ambient oxygen partial pressure. Different letters indicate significant (*P* < 0.05, Tukey–Kramer test) differences across the treatments. *P*-values < 0.05 [from one sample t-test] indicate a significant shift of V_cmax_ and J_max_ (from a 1:1 ratio, cf. Fig. 6) due to changes of partial pressure of oxygen in air. Treatments: WC = Well-watered control, WP = Well-watered phloem cut, WX = Well-watered xylem cut, WXP = Well-watered xylem-phloem cut, DC = Drought control, DP = Drought phloem cut, DX = Drought xylem cut, DXP = Drought xylem-phloem cut.

| Treatments | $V_{\text{cmax}}$<br>[Ambient O<sub>2</sub>] | $J_{\text{max}}$<br>[Ambient O<sub>2</sub>] | $V_{\text{cmax}}$ | | $J_{\text{max}}$ | |
| --- | --- | --- | --- | --- | --- | --- |
| | | | Ratio<br>[Low/<br>Ambient O<sub>2</sub>] | $P$ value | Ratio<br>[Low/<br>Ambient O<sub>2</sub>] | $P$ value |
| WC | 104.2±3.6 A | 148.1±4.1 A | 1.09±0.03 | 0.07 | 0.96±0.03 | 0.29 |
| WP | 91.2±5.1 ABC | 127.4±9.4 A | 0.88±0.04 | **0.01** | 0.79±0.06 | **0.03** |
| WX | 76.5±10.3 BC | 123.2±18.7 A | 0.97±0.11 | 0.85 | 0.94±0.05 | 0.38 |
| WXP | 87.5±9.3 ABC | 127.3±15.3 A | 1.05±0.12 | 0.66 | 1.02±0.10 | 0.78 |
| DC | 85.5±1.9 ABC | 132.1±4.1 A | 0.96±0.06 | 0.67 | 0.95±0.08 | 0.62 |
| DP | 99.6±4.8 AB | 136.4±14.7 A | 0.88±0.07 | 0.16 | 0.93±0.16 | 0.71 |
| DX | 69.2±8.1 C | 126.7±7.7 A | 0.99±0.14 | 0.97 | 0.90±0.03 | 0.08 |
| DXP | 79.3±10.7 ABC | 113.1±7.8 A | 0.93±0.10 | 0.62 | 0.98±0.12 | 0.87 |

### Non-structural carbohydrates

Overall, total non-structural carbohydrate concentrations (NSC) were significantly affected by the drought treatment and by the cut treatments, the latter mostly in leaves and in particular when phloem damage was involved. Droughted plants had higher total NSC concentration in stems and roots, and marginally higher ones in leaves compared to well-watered plants (Fig. 5; Table S1), and this was always due to increased soluble sugars concentrations. Within the droughted treatments, NSC concentrations of leaves, stems or roots did not differ significantly among the cut treatments (Fig. 5), although trends in leaves of droughted plants were similar to those in well-watered plants. In well-watered plants, NSC concentrations of leaves were increased in plants that included a phloem cut, and these increases were largely due to an increase in starch. Leaf starch concentrations were 84% in WP and 75% in WXP relative to those in well-watered control plants (WC), and were 73% in DP and 12% in DXP relative to those in droughted control plants (DC). Similar but less pronounced patterns were observed in stems, whereas opposite patterns occurred in roots, where starch and NSC were lower in plants that had experienced a phloem cut. Root starch concentrations were 51% in WP and 71% in WXP relative to the control (WC). Root starch concentrations were uniformly low in roots of droughted plants. Across all the treatments, photosynthesic rate was negatively correlated with leaf soluble sugar concentration (droughted: R^2^ = 0.37; P = 0.003; well-watered: R^2^ = 0.32; P = 0.005; Fig. S4).

**Figure 5:**
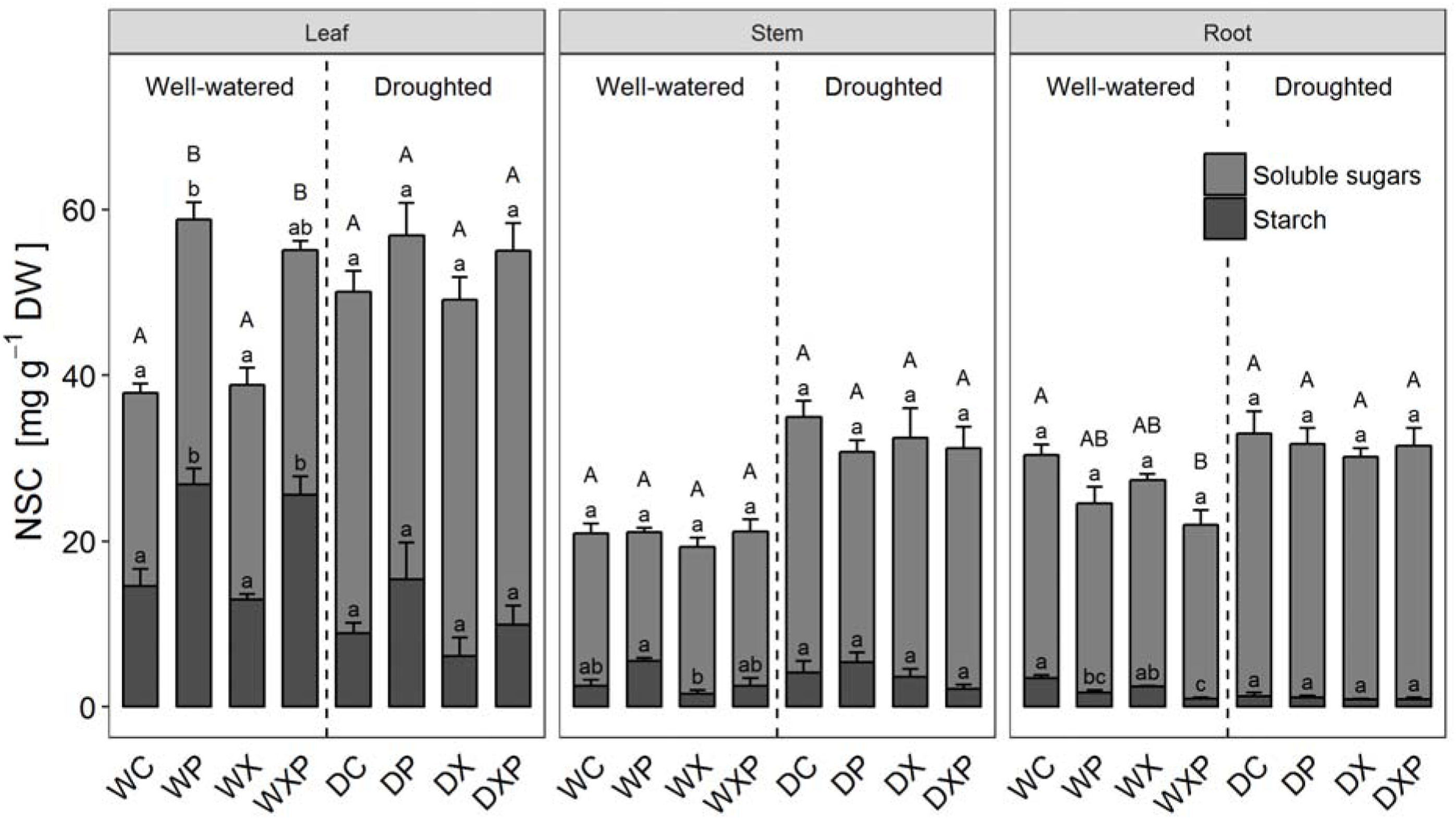
Non-structural carbohydrates (NSC) in leaf, stem and root of the plants under different treatment at harvest. Different capital letters indicate significant (*P* < 0.05, Tukey–Kramer test) difference of NSC and smaller letters indicate significant difference of soluble sugars and starch between cut treatments within each water treatment. Treatments: WC = Well-watered control, WXC = Well-watered xylem control, WP = Well-watered phloem cut, WX = Well-watered xylem cut, WXP = Well-watered xylem-phloem cut, DC = Drought control, DXC = Drought xylem control, DP = Drought phloem cut, DX = Drought xylem cut, DXP = Drought xylem-phloem cut.

## Discussion

Our experiment tested the effects of severe damage to xylem and phloem, similar to that caused by a stem canker, and the potential exacerbation of such effects by drought stress. Interestingly, the impact of stem damage was greater in well-watered plants than drought-stressed plants. As we hypothesized, xylem damage reduced whole-plant conductance, leaf stomatal conductance and photosynthesis, and hence growth. Phloem damage interrupted phloem transport, causing some changes in non-structural carbohydrate concentrations in leaves and roots, and reducing photosynthesis. Plants were remarkably resilient to significant loss of xylem and phloem, although xylem damage (55% loss tissue, on average) had a relatively greater impact on most of the physiological parameters than phloem damage (70% loss of tissue, on average). The effect of combined damage to both xylem and phloem on plant physiological responses was never more (and often less) than additive.

In contrast to our hypothesis, we found no evidence of greater impact of drought stress in plants with xylem damage compared to undamaged controls. Loss of stem xylem did not significantly reduce whole-plant hydraulic conductance in drought-stressed plants, unlike in well-watered plants. Whilst it did reduce stomatal conductance, and therefore transpiration, the percentage reduction was similar in drought-stressed and well-watered plants. The combined result of physiological behaviour and morphological/anatomical adjustments was that midday water potentials were kept within a narrow range in all drought treatments, just below -2 MPa. Such water potentials are commonly observed in this species in glasshouse and field studies (Poot & Veneklaas, 2013; Hossain *et al*., 2018), and the fact that xylem-damaged plants did not have lower water potentials than non-damaged plants indicates that plants managed to avoid critical levels of dehydration through stomatal regulation (Szota *et al*., 2011). The morphological/anatomical adjustments may have included a reduction in the ratio of leaf area to sapwood area, and an increase in sapwood specific conductivity (greater proportion of xylem vessels functional; more and wider xylem vessels). The remarkable resilience to experimental removal of more than half of plants’ sapwood confirms observations on stem canker-affected plants of this same species (Hossain *et al*., 2018). The observation in the current experiment that xylem damage affected growth and physiology of well-watered plants more than of droughted plants, may be due to a greater reliance on xylem function when plants are growing faster and using more resources. For example, the higher transpiration rate of well-watered plants may require the xylem to function at full capacity, whereas xylem transport may not limit transpiration rate of droughted plants.

For phloem damage treatments we had hypothesised that well-watered plants would be more affected than drought-stressed plants, as we expected that carbon transport would be less critical in stressed plants that have lower levels of metabolism. Our results do indeed show larger effects in well-watered plants. Whilst the negative effects of phloem damage on plant function were generally smaller than the effects of xylem damage, they were statistically significant. Effects of phloem damage presumably acted through impeded sugar transport, leading to increased NSC in leaves and decreased NSC in roots (Asao & Ryan, 2015; Mitchell *et al*., 2017; Rainer-Lethaus & Oberhuber, 2018). These changes were relatively small, and statistical significance was largely limited to starch fractions in well-watered plants. We hypothesise that the modest changes in NSC pools despite substantial transport impediments can be explained by increased sugar transport through the remaining phloem connections (i.e. redundancy of transport capacity in intact plants), and adjustments in rates of photosynthesis and respiration. Our measurements of photosynthetic parameters are indicative of feedback inhibition of photosynthesis (see below). The mechanism of growth depression, however, is unknown: it is clear that there was no carbon limitation in leaves of phloem-damaged plants, and therefore a reduced sink activity was likely responsible (Körner, 2015). In the absence of environmental stress (except for the drought treatment, but drought stress was not exacerbated by phloem damage) hormonal signals were likely implicated. No significant effects of phloem damage on plant water relations were detected, other than reduced stomatal conductance. Reduced stomatal conductance has also been documented for girdled *Pinus canariensis* by López *et al*. (2015), who attributed it to feedback inhibition of photosynthesis and hormonal (abscisic acid) effects. In addition, reduced photosynthate transport due to phloem damage may hinder the ability to reverse xylem embolisms, and thus may indirectly reduce stem conductivity and stomatal conductance (Lachenbruch & Zhao, 2019). Phloem damage may also reduce root growth and thus cause a lower root/shoot ratio, which could limit water supply to foliage. While there was a tendency for root/shoot ratio to be lowest in phloem-damaged plants, this is unlikely to have affected stomatal conductance, as the differences were not statistically significant, and were only apparent in well-watered plants where water limitation is not expected.

Feedback inhibition of photosynthesis in our experiment is supported by two lines of evidence: 1. Negative correlations between photosynthetic rate and leaf soluble sugar concentrations (Fig. S3); 2. Absence of a positive effect of low O_2_ on V_cmax_ and J_max_ (Table 1). This oxygen insensitivity of photosynthesis (or reverse sensitivity: V_cmax_ and J_max_ were significantly lower at low O_2_ in well-watered-phloem-damaged plants) is interpreted as feedback inhibition of photosynthesis (Morcuende *et al*., 1997; Sun *et al*., 1999), presumably caused by accumulation of Calvin Cycle intermediates (triose phosphates), limiting the recycling of Pi back to chloroplasts and reducing photosynthesis (Lambers *et al*., 2008; Ellsworth *et al*., 2015). The increased starch formation, which we observed in phloem-damaged plants, counteracts Pi sequestration to some extent. It must be noted that photosynthesis of plants without phloem or xylem damage also did not show a significant positive response to low O_2_, suggesting that they may also have been sink-limited to some extent, or that photorespiration was of relatively minor importance.

The experiment allowed us to assess potential interactive effects of combined phloem and xylem damage. Given the inherent interdependence of the two transport systems (De Schepper *et al*., 2013; Sevanto, 2014; Perri *et al*., 2019), it was expected that the considerable levels of damage to both systems would have greater than additive effects on growth, water status and carbohydrates. Although growth was lowest in the combined phloem/xylem damage treatment, it was not significantly lower than in the treatment that had xylem damage only. Plant water status and hydraulic parameters of phloem/xylem-damaged plants were not different from plants with xylem damaged only, and photosynthetic and carbohydrate parameters were not different from plants with phloem damage only. It appears therefore that the observed effects on plant function could be explained by the impacts on the transport systems most directly associated with the respective functions. It is important, however, to stress that plants with damage to both phloem and xylem showed all the impacts that were observed for plants that had damage to either phloem or xylem, and were therefore most severely compromised. Such plants are comparable to plants affected by the stem canker *Quambalaria coyrecup*.

Our experiment demonstrates the considerable resilience of *C. calophylla* subject to very substantial loss of xylem and phloem over a period of four months, both in well-watered and water-limited conditions. Reduced growth and water use, morphological and physiological adjustments, and presumably a degree of redundancy of transport capacity in healthy plants prevented serious negative consequences of the treatments. It is to be expected, however, that there are limits to the ability of trees to adjust to stem damage, for example due to stem cankers. Whilst our experimental treatments simulated severe stem damage, the damage was not progressive, unlike the situation in many canker-affected stems, where growth of new stem tissue is the only way to keep ahead of tissue loss through disease. Even if damage is not progressive, certain consequences of xylem and phloem damage may not manifest themselves until more time has passed. For example lack of growth may only become fatal when naturally senescing tissues, including leaves, cannot be replaced. The probability of experiencing more severe stress, for example, extreme temperature events or long-term drought, will also increase with time. The inability to repair xylem embolism or replace damaged leaves could prove fatal. Future research should assess these longer-term and higher-intensity stress factors.

## Acknowledgments

We acknowledge Trudy Paap for providing with the seedlings used in this experiment. Funding was provided by the Australian Research Council (Linkage Project 120200581), and School of Plant Biology, University of Western Australia. We also thank the reviewers for their helpful suggestions.

## Author Contributions

M.H. led the concept development, designed and conducted the research, analysis, and wrote the paper. E.V and P.P. contributed to the concept development and experimental design, and edited the manuscript.

## Supporting Information

**Table S1.** Summary of statistical analysis for various physiological parameters as dependent on watering and cut treatments, and their interactions. Values in the table are P values from ANOVA.

| Parameters | Cut treatment | Drought treatment | Cut * Drought treatment |
| --- | --- | --- | --- |
| RGR of total stem length | **<0.01** | **<0.001** | **<0.001** |
| RGR of diameter above cut | **0.02** | **<0.001** | **<0.001** |
| RGR of diameter below cut | 0.38 | **<0.001** | **<0.001** |
| Whole plant dry biomass | **<0.001** | **<0.001** | **<0.001** |
| Root:shoot ratio | 0.63 | **<0.001** | **0.004** |
| Leaf area to sapwood area | **<0.001** | 0.29 | **0.04** |
| leaf area to total plant dry mass<br>(Leaf area ratio) | 0.94 | **0.01** | 0.81 |
| Predawn WP | 0.95 | **<0.001** | **<0.001** |
| Midday WP | 0.38 | **<0.001** | **<0.001** |
| K<sub>p</sub> | **<0.001** | **<0.001** | **<0.001** |
| A<sub>sat</sub> | **<0.001** | **<0.001** | **<0.001** |
| g<sub>s</sub> | **<0.001** | **<0.001** | **<0.001** |
| V<sub>cmax</sub> (ambient O<sub>2</sub>) | 0.15 | 0.34 | 0.33 |
| J<sub>max</sub> (ambient O<sub>2</sub>) | 0.27 | 0.38 | 0.55 |
| TPU (ambient O<sub>2</sub>) | 0.71 | 0.14 | 0.40 |
| V<sub>cmax</sub> (low O<sub>2</sub>) | **0.003** | 0.28 | 0.15 |
| J<sub>max</sub> (low O<sub>2</sub>) | **0.01** | 0.85 | 0.62 |
| TPU (low O<sub>2</sub>) | 0.11 | 0.31 | 0.51 |
| Leaf NSC | **0.007** | **<0.001** | **<0.001** |
| Leaf soluble sugars | 0.34 | **<0.001** | **<0.001** |
| Leaf starch | **<0.001** | **<0.001** | **<0.001** |
| Stem NSC | 0.78 | **<0.001** | **<0.001** |
| Stem soluble sugars | 0.59 | **<0.001** | **<0.001** |
| Stem starch | **0.04** | **0.12** | **0.04** |
| Root NSC | 0.14 | **<0.001** | **0.006** |
| Root soluble sugars | 0.51 | **<0.001** | **0.001** |
| Root starch | **<0.001** | **<0.001** | **<0.001** |

**Figure S1.**
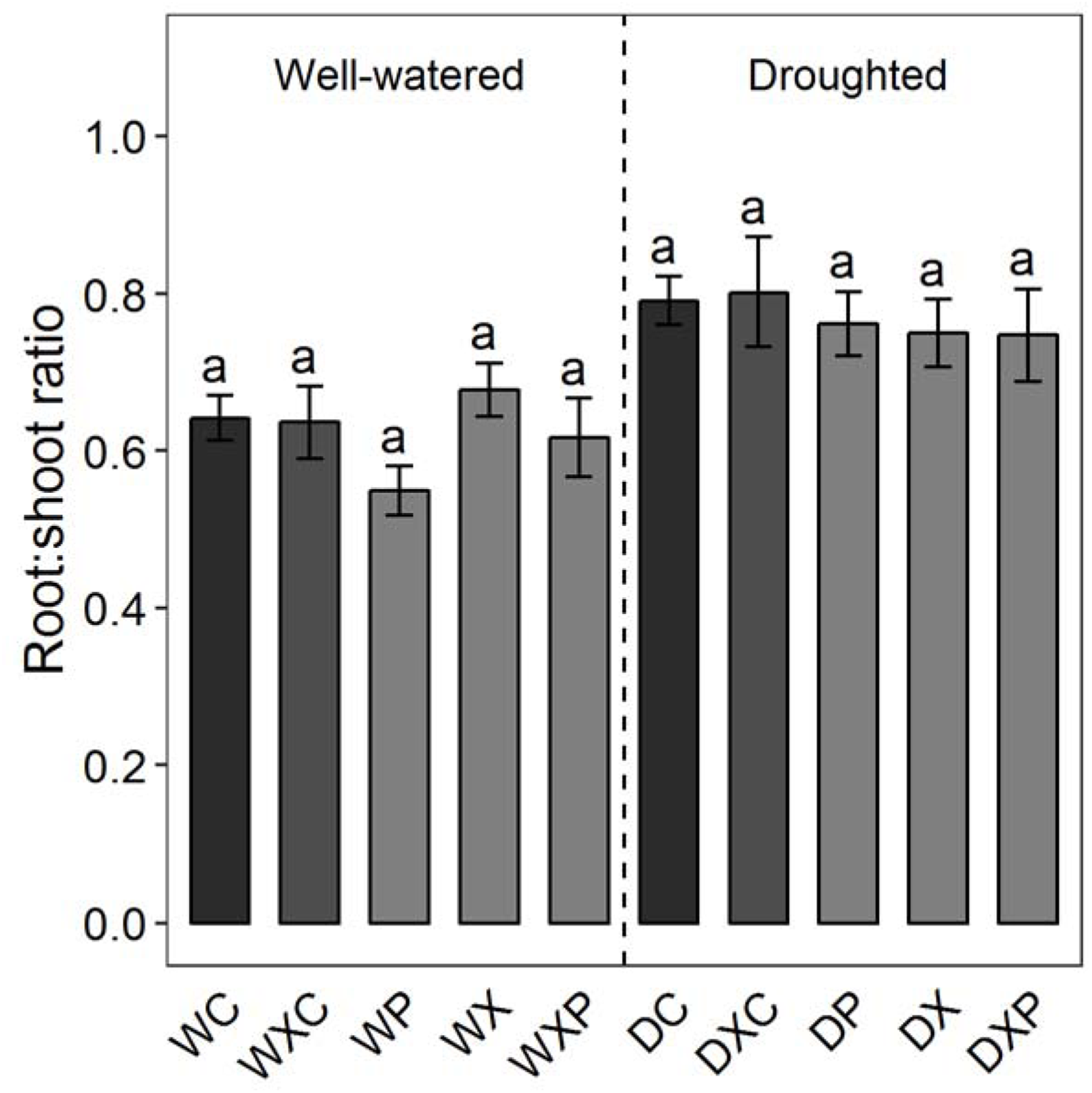
Root: shoot ratio of the plants under different treatments at harvest. Different letters indicate significant (P < 0.05, Tukey–Kramer test) differences between cut treatments within each water treatment. Treatments: WC = Well-watered control, WXC = Well-watered xylem control, WP = Well-watered phloem cut, WX = Well-watered xylem cut, WXP = Well-watered xylem-phloem cut, DC = Drought control, DXC = Drought xylem control, DP = Drought phloem cut, DX = Drought xylem cut, DXP = Drought xylem-phloem cut.

**Figure S2.**
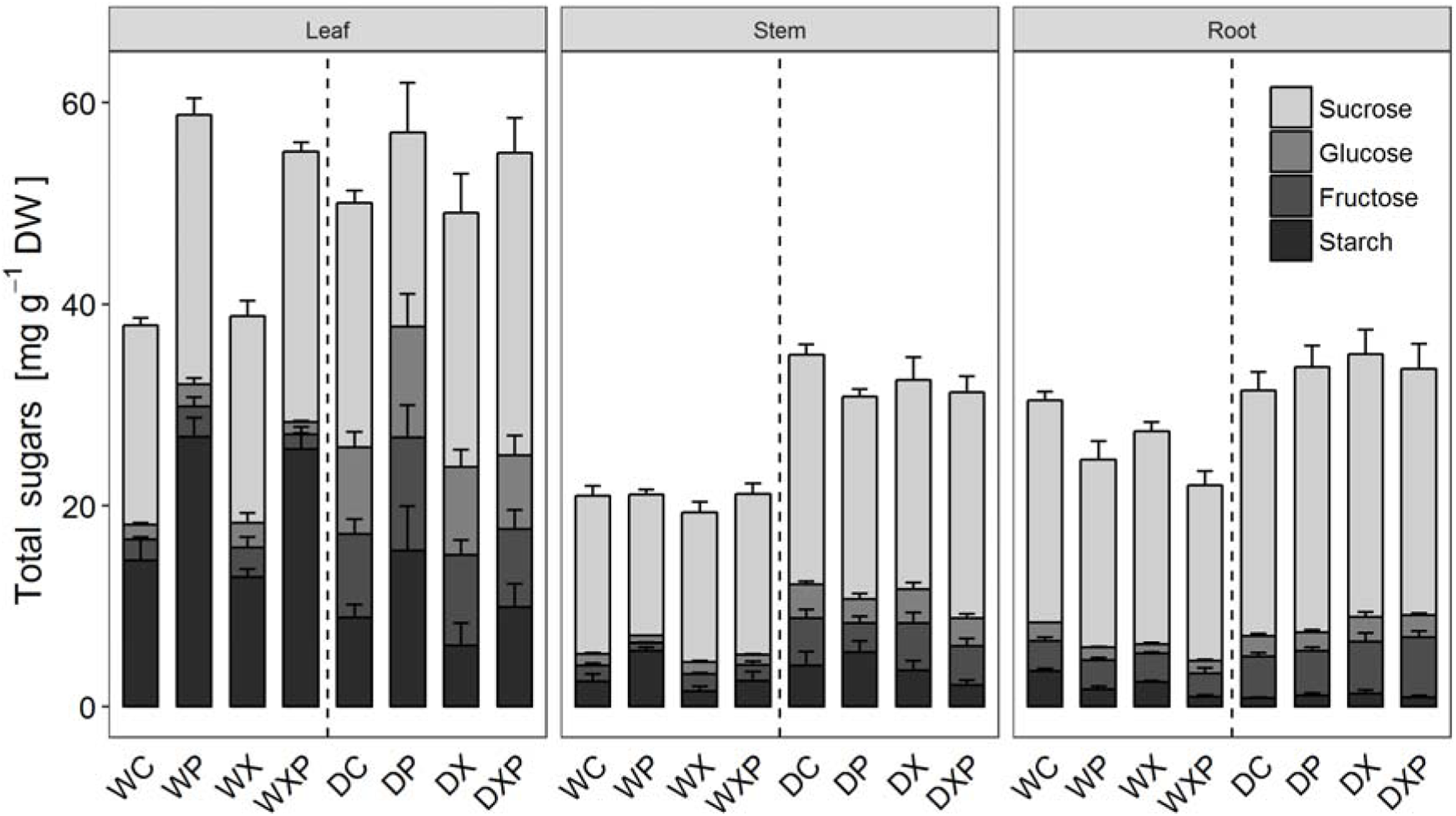
NSC represented by the average sucrose, fructose and glucose and starch concentrations in leaves, stems and roots of the plants under different treatments at harvest. Treatments: WC = Well-watered control, WP = Well-watered phloem cut, WX = Well-watered xylem cut, WXP = Well-watered xylem-phloem cut, DC = Drought control, DP = Drought phloem cut, DX = Drought xylem cut, DXP = Drought xylem-phloem cut.

**Figure S3.**
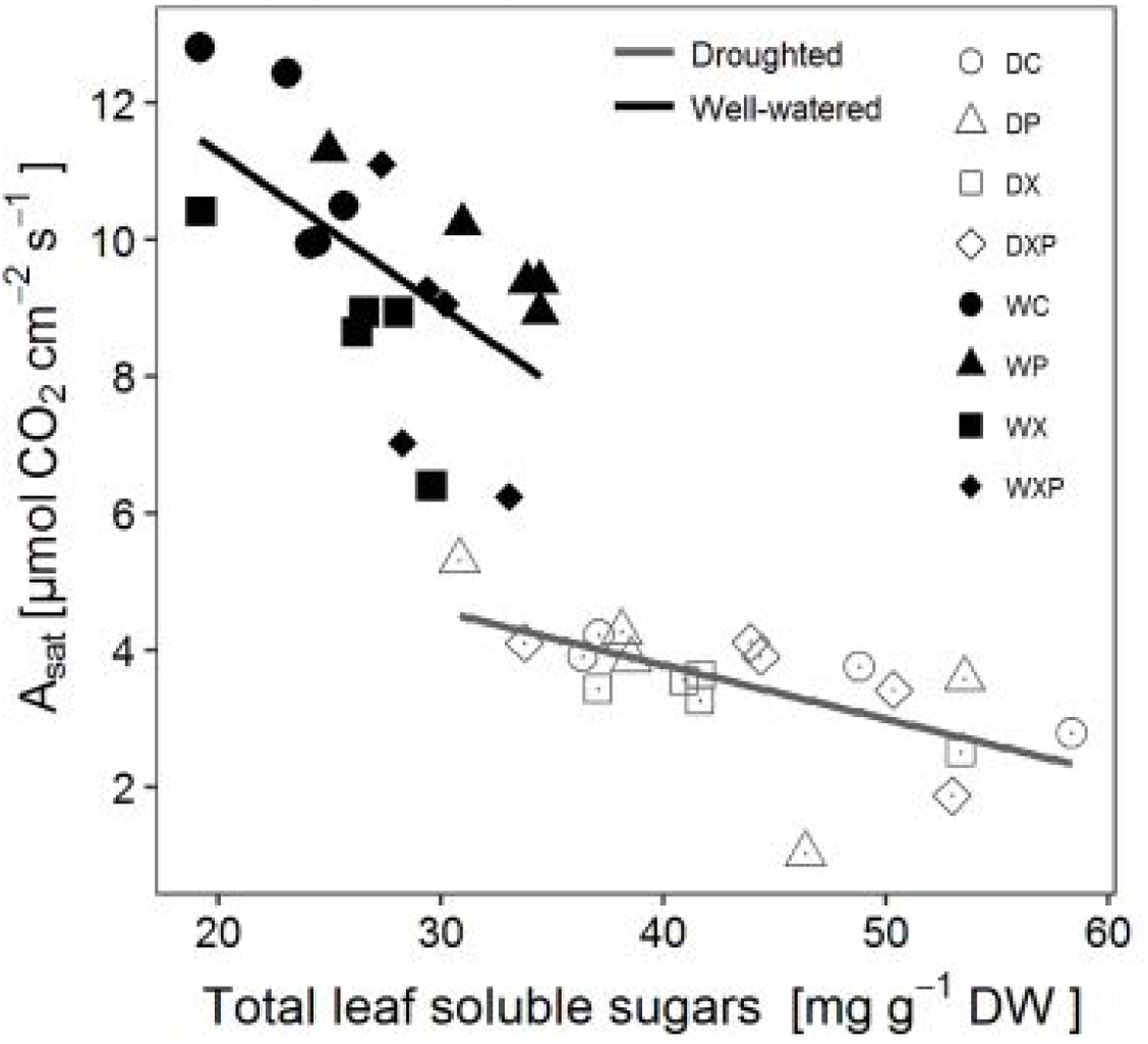
Relationship between net photosynthetic rate at saturating light intensity (A_sat_) and total leaf soluble sugar concentration for the plants under different water and cut treatments. Data are fitted with a linear model (well-watered: y = -0.23x + 15.82, R = 0.32, P = 0.005; and droughted: y = -0.07x + 6.93, R = 0.37, P = 0.003). Treatments: WC = Well-watered control, WP = Well-watered phloem cut, WX = Well-watered xylem cut, WXP = Well-watered xylem-phloem cut, DC = Drought control, DP = Drought phloem cut, DX = Drought xylem cut, DXP = Drought xylem-phloem cut.

